# A sucrose non-fermenting-1-related protein kinase 1 gene from wheat, TaSnRK1α regulates starch biosynthesis by modulating AGPase activity

**DOI:** 10.1101/2023.07.01.547320

**Authors:** Prashant Kumar, Akansha Madhawan, Akshya Sharma, Vinita Sharma, Deepak Das, Afsana Parveen, Vikas Fandade, Deepak Sharma, Joy Roy

## Abstract

Major portion of wheat grain consist of carbohydrate, mainly starch. The proportion of amylose and amylopectin in starch greatly influence the end product quality. Advancement in understanding starch biosynthesis pathway and modulating key genes has enabled the genetic modification of crops resulting in enhanced starch quality. However, the regulation of starch biosynthesis genes still remains unexplored. So, to expand the limited knowledge, here, we characterized a Ser/Thr kinase, SnRK1α in wheat and determined its role in regulating starch biosynthesis. SnRK1 is an evolutionary conserved protein kinase and share homology to yeast SNF1. Yeast complementation assay suggest *TaSnRK1*α restore growth defect and promotes glycogen accumulation. Domain analysis and complementation assay with truncated proteins suggest the importance of ATP-binding and UBA domain in TaSnRK1α activity. Sub-cellular localization identified nuclear and cytoplasmic localization of TaSnRK1α in tobacco leaves. Further, heterologous over-expression (O/E) of *TaSnRK1*α in Arabidopsis not only led to increase in starch content but also enlarges the starch granules. *TaSnRK1*α was found to restore starch accumulation in Arabidopsis *kin10.* Remarkably, *TaSnRK1*α O/E increase the AGPase activity suggesting the direct regulation of rate limiting enzyme AGPase involved in starch biosynthesis. Furthermore, in vitro and in vivo interaction assay reveal that TaSnRK1α interacts with AGPase large sub-unit. Overall, our findings indicate that TaSnRK1α plays role in starch biosynthesis by regulating AGPase activity.

**Highlights:**

- TaSnRK1α is Ser/Thr kinase in wheat and show dual localization in nucleus and cytoplasm.
- Overexpression of TaSnRK1α increases starch content and enlarges starch granules in Arabidopsis.
- TaSnRK1α enhances AGPase activity thereby regulating starch biosynthesis.
- TaSnRK1α directly interact with AGPase large subunit in vivo and in vitro.

## Introduction

Bread wheat (*Triticum aestivum* L.) is a major staple food crop and a key source of calories intake and nutrition for human population. The wheat grain consists of approximately 70 % carbohydrate, 13 % water and small portion of proteins and lipids. Starch is the major carbohydrate of the wheat grain (Shewry and Halford, 2002) and constitutes a substantial portion of the grain (Kim and Kim, 2021). Wheat starch mainly consist of two polymers called amylose and amylopectin (Nakamura *et al*., 1995; Tester *et al*., 2004). In natural conditions, amylose constitutes 25-30 % while amylopectin constitutes 70-75 % of the total starch while both have different structure and properties (Alcázar-Alay and Meireles, 2015; Chen *et al*., 2016). Amylose is a linear polymer linked by α-1,4 glycosidic bonds whereas amylopectin is more branched structure linearly linked by α-1,4 glycosidic bonds and α-1,6 glycosidic bonds at branching points (French *et al*., 1984; Tetlow and Bertoft, 2020). Amylose and amylopectin ratio in starch, is a key factor to determine and influence the starch quality and its physiochemical properties.

Starch biosynthesis in wheat endosperm involves coordinated activities of different enzymes, each with its specific function. The first committed and rate limiting step in starch biosynthesis is AGPase reaction, which converts glucose-1-phosphate and ATP to ADP- glucose (ADPG), which serves as a substrate for starch synthases (Jeon *et al*., 2010). Soluble starch synthases (SS) work on ADPG and are mainly responsible for amylopectin biosynthesis along with branching and debranching enzymes (Keeling and Myers, 2010; Goren *et al*., 2018; Sharma *et al*., 2022). There are different isoforms of SS with specific roles; SSI is responsible for elongating the short glucan chains with degree of polymerization (DP) of 6-10 units while SSII elongates the longer glucan chains with a DP of 12-20 units. SSIII has been implicated in the synthesis of even longer glucan chains while SSIV is involved in starch granules initiation (Delvallé *et al*., 2005; Roldán *et al*., 2007; Zhang *et al*., 2008). Starch branching and debranching enzymes along with starch phosphorylase are responsible for formation and positioning of branching in amylopectin and the removal of excess branches wherever necessary (Tetlow and Emes, 2014). GBSSI is primarily expressed in the amyloplast of cereal endosperm and is the key enzyme of amylose biosynthesis (Nishi *et al*., 2001; Nakamura *et al*., 2006; Regina *et al*., 2006). It plays a crucial role in starch biosynthesis by elongating linear glucose chains using ADPG as the substrate. Another isoform of GBSS, GBSSII is involved in amylose biosynthesis in leaves, contributing to transitory starch. The key pathway of starch biosynthesis is well known but the regulatory mechanism is not well studied in cereal crops including wheat. The regulatory factors associated with starch biosynthesis and their roles in improving starch quality are not explored in wheat.

Kinases are one of the key modulators of different metabolic pathways and their activities underpin many basic cellular processes. The plant sucrose non-fermenting-1 (SNF1)-related kinase 1 (SnRK1), animal AMP-activated protein kinase (AMPK), and yeast SNF1 kinase are members of evolutionarily conserved serine/threonine protein kinases (Polge and Thomas, 2007). These kinases play crucial role in energy homeostasis, cellular responses to nutrient availability, and control of carbohydrate metabolism. SnRK1 and its orthologs SNF-1 and AMPK, consist of a catalytic α sub-unit and regulatory sub-units β and γ in plants (Peixoto and Baena-González, 2022). There are various studies, where the role of SnRK1 in starch biosynthesis is reported. Zhang *et al*., (2001) reported loss of starch accumulation in pollen grains of barley by expressing the antisense SnRK1 sequence and Kanegae *et al*., (2005) found accumulation of high starch by SnRK1 overexpression in rice caryopsis. Moreover, an increase of about 30% in starch accumulation was observed by overexpressing the SnRK1 in potato tubers (McKibbin *et al*., 2006). Several other studies also suggested the involvement of SnRK1 in high starch biosynthesis such as in sweet potato (Ren *et al*., 2019) and transgenic tomato (Wang *et al*., 2012). Above studies demonstrated that overexpression of SnRK1 is responsible for upregulation of expression and activity of key enzymes. Jiang *et al*., (2013) reported increased activity of SuSy and AGPase by overexpression of SnRK1 in tobacco, whereas Wang *et al*., (2017) reported increased activity of AGPase and SIII in tobacco by overexpression of *StSnRK1*. Ren *et al*., (2019) found 7 fold increased activity of AGPase by overexpression of *IbSnRK1* in sweet potato. However, the exact mechanism by which SnRK1 regulates starch biosynthesis still remains unclear.

In our previous study, we identified TaSnRK1α transcriptionally up-regulated in three high amylopectin EMS induced wheat mutant lines (Kumar *et al*., 2022). In this study, we systematically characterized TaSnRK1α and its role in regulation of wheat starch biosynthesis. Using the yeast system, we demonstrated the role of TaSnRK1α in glycogen accumulation. Further, we reported that total starch content is increased when *TaSnRK1*α is overexpressed in *Arabidopsis thaliana* ecotype Col-0 and *Nicotiana benthamiana*. Our results also demonstrated that TaSnRK1α can restore starch accumulation in *kin10* Arabidopsis mutant lines. Sub-cellular localization revealed the dual localization of TaSnRK1α in nucleus as well as cytoplasm. Finally, we explored the underlying mechanism of TaSnRK1α, using in vivo and in vitro studies. Our results revealed that TaSnRK1α interacts with AGPase large subunit, thereby, modulating the AGPase activity and hence we concluded that TaSnRK1α, a Ser/Thr kinase, could play an important role in starch biosynthesis pathway in wheat.

## Materials and Methods

### In silico analysis of wheat SnRK1

Wheat SnRK1α amino acid and nucleotide sequences were retrieved from the Ensembl Plants database (https://plants.ensembl.org/index.html). Domain analysis of TaSnRK1α was carried out using the amino acid sequence, by InterProScan (https://www.ebi.ac.uk/interpro/), UniProt (https://www.uniprot.org/), SMART (http://smart.embl-heidelberg.de/) and CDD (https://www.ncbi.nlm.nih.gov/Structure/cdd/cdd.shtml) web tools and databases.

For phylogenetic analysis, amino acid sequences of various cereals (Barley, maize, rice, rye, and sorghum) along with Arabidopsis, and other dicot plants having rich starch content (Banana, Sweet potato, and Potato) were retrieved using the BlastP. The sequence alignment of the amino acid sequences was performed using the ClustalW (https://www.ebi.ac.uk/Tools/msa/clustalw2/) and a phylogenetic tree was constructed using the MEGAX software (https://www.megasoftware.net/). Maximum-likelihood method was used for phylogeny analysis with 1000 bootstraps per replication. Further, Interactive Tree of Life (IToLv6, http://itol.embl.de) was used for phylogenetic tree display.

For T-loop activation domain identification, amino acid sequences of Arabidopsis SnRK1 (*KIN10*), human AMPK, *Hordeum vulgare* SnRK1, rice SnRK1, wheat SnRK1 and maize SnRK1 were retrieved from Ensembl Plants and aligned using the online web-server tool T- Coffee (https://www.ebi.ac.uk/Tools/msa/tcoffee/).

### Recombinant protein expression and kinase assay

The enzymatic activity of the TaSnRK1α was assessed using the Universal Fluorometric Kinase Assay Kit (Sigma). For kinase assay, the CDS sequence of *TaSnRK1*α was cloned into the protein expression vector pET28a+ and further transformed into BL21 (DE3) cells. Primer details are provided in Supplementary Table 1. His-tagged recombinant protein was purified using Ni-NTA purification with the help of Ni-charged resins (Bio-Rad). Purified protein was further used for kinase assay. A total of 5 μM of the enzyme was used in the kinase assay mix that contains 12.5 μM AMARA peptide, a substrate for SnRK1, and 25 µM ATP. This mixture was incubated at 37 °C for 30 mins and recorded at an excitation wavelength of 540 nm and emission wavelength of 590 nm. All the assays were done in triplicates.

### Yeast complementation assay

To elucidate the role of wheat SnRK1 in carbohydrate metabolism in yeast, the full length CDS of *TaSnRK1*α was cloned in pYES2 yeast expression vector using restriction enzyme sites BamHI and XhoI. Primer details are provided in Supplementary Table 1. Empty vector and TaSnRK1α were transformed into the mutant strain Δ*snf1* and wild type strain CEN.PK using the Li-Ac protocol (Gietz and Schiestl, 2007). Mutant strain Δ*snf1* was kindly provided by Dr. Sunil Laxman (InStem, Bengaluru, India). An empty vector was used as a control. The incompetency of yeast mutant Δ*snf1* to grow on sucrose media was used as an assay.

For the assay, cultures of positive transformants were grown at 30 °C on minimal synthetic media without uracil (SD/-Ura) with 2% glucose, and complementation assay was performed on SD/-Ura with 2% glucose or 2% sucrose as different carbohydrate source media, with different dilutions. 10 µl of each dilution was plated on the SD/-Ura plates and growth was analysed after 3 days at 30 °C.

### Glycogen estimation in yeast cells

Further, to explore the role of TaSnRK1α in glycogen accumulation, glycogen was estimated qualitatively and quantitatively. For histochemical studies, cells from each yeast transformant were plated on a SC-gal plates and incubated at 30 °C for 24 hours. Plated colonies were stained with Lugol’s iodine solution and imaged immediately. For light microscopy, liquid cultures from each transformant were grown in complex media (containing 2% galactose and 1% raffinose) and stained in one volume of Lugol’s solution and imaged under automated Leica DM6000 B and images were acquired under 20X and 40X magnification. For quantitative estimation, yeast primary cultures from each transformant were grown overnight at 30 °C with shaking at 180 RPM. Primary cultures were used to inoculate the secondary culture of 750 ml per replicate and grown at 30 °C for 48 hours. Cells were pelleted for each strain and glycogen was extracted by KOH mediated alkali method previously described by Aklujkar *et al*., (2008). Briefly, cells were pelleted by centrifugation then washed twice with distilled water and treated with 20 % KOH per gm of the dry weight of the cells. Treated cells were boiled for 1 hour, then the pH was adjusted to 7 using HCl. After centrifugation, the insoluble fraction and the soluble fraction were separated. Both the fractions were digested to glucose by treating with α-amylase and amyloglucosidase (Megagyme), at 37 °C for 6-7 hours and the total liberated glucose was calculated using the DNSA method (Rajbhar *et al*., 2015).

To characterize the glycogen extracted from yeast, FTIR spectra was performed using the infrared spectrometer 536 (ATR-FTIR) (PerkinElmer UATR Spectrum Two). Spectra of glycogen were recorded between 2000 to 500 cm^-1^. Background spectra of a clean ATR surface were acquired prior to each sample measurement using the same acquisition parameters. Each sample were scanned three times to observe good reproducibility.

### Domain deletion and activity in yeast

To study the effect of the UBA and ATP binding domain on the function of TaSnRK1α, UBA and ATP binding domains were deleted using the PCR-based plasmid deletion approach. Both the domain deleted truncated TaSnRK1α genes were cloned individually into the yeast expression vector pYES.2 using BamHI and XhoI as restriction enzyme sites. Primer details are provided in Supplementary Table 1. Clones were confirmed through restriction enzyme digestion and sequencing. The effect of domain deletion on TaSnRK1α was assessed using the yeast complementation assay by observing the growth defect on different carbohydrate sources in media.

### Sub-cellular localization of wheat SnRK1

In silico sub-cellular localization of the TaSnRK1α was predicted using the online web-server BUSCA (https://busca.biocomp.unibo.it/). To analyse, the sub-cellular localization of wheat SnRK1, the full length CDS of *TaSnRK1*α was cloned into the plant expression vector pCAMBIA1302 without stop codon under 35S promoter, with GFP protein fused at C-terminal and NcoI restriction enzyme sites on both ends using Gibson Assembly (NEB, USA). Empty vector pCAMBIA 1302 was used as a positive control. Primer details are provided in Supplementary Table 1.

The plasmids carrying 35S:*TaSnRK1*α:GFP and empty vector were transformed into *Agrobacterium tumefaciens* GV1301 strain using the freeze-thaw method (Chen *et al*., 1994). To explore the localization, primary cultures of 35S:*TaSnRK1*α:GFP and empty vector were grown overnight at 28 °C for secondary culture. Secondary cultures were grown till O.D. reaches 0.8-1. The secondary culture was pelleted and resuspended into the resuspension buffer containing MES- 10 mM, MgCl_2_- 10 mM, and acetosyringone- 200 μM, incubated at room temperature for 4-5 hours. Resuspension solution was infiltrated in young leaves of 4-5 weeks old tobacco (*N. benthamiana*) plants. Plants were kept in dark for 24 hours after infiltration and then shifted to growth chamber at 22 °C for 3 days. Infected leaves were seen under laser scanning confocal microscopy on Carl Zeiss confocal microscope LSM880 (Germany).

An immunoblot assay was performed to further confirm the 35S:TaSnRK1α:GFP tagged fusion protein expression in the tobacco leaves using wheat specific anti-TaSnRK1α specific antibody with 1:5000 dilution using the wet-transfer buffer system.

### Heterologous expression in Tobacco and starch estimation

To further gain insights into the role of TaSnRK1α in starch biosynthesis in planta, full length CDS of *TaSnRK1*α gene was amplified using the wheat cDNA and cloned into the pCAMBIA1302 vector under the constitutive promoter 35S using the Gibson assembly cloning kit (Gibson Assembly® Master Mix, NEB) with the help of restriction enzyme NcoI. Primer details are provided in Supplementary Table 1. Recombinant plasmid was further transformed into the *Agrobacterium tumefaciens* GV1301 strain with appropriate antibiotics. 35S:TaSnRK1α construct along with empty vector was infiltrated into the fresh, healthy leaves of 4 weeks old tobacco plants. After a week, starch estimation was performed using the previously described enzymatic reaction with the help of α-amylase and amyloglucosidase, and liberated glucose was estimated using the DNSA reagent.

### Arabidopsis *TaSnRK1*α overexpression lines

To see the effect of *TaSnRK1*α overexpression on Arabidopsis, full length CDS of TaSnRK1α was cloned into plant expression vector pCAMBIA1302 under the constitutive promoter CaMV:35S with the help of restriction site NcoI on both ends using Gibson assembly® master mix (NEB, USA). Primer details are provided in Supplementary Table 1. 35S:TaSnRK1α along with empty vector, were transformed into Agrobacterium cells (GV1301) and used for plant transformation. For plant transformation, wild type ecotype Col- 0 was grown in a plant growth chamber at 22 °C, with 16 h light and 8 h dark period till the flowering stage. Plants were transformed using floral dip method (Zhang *et al*., 2006) with above Agrobacterium construct. Seeds were collected after maturation and multiple transformants were selected on half MS plates under hygromycin selection. Positive healthy plants with green and long hypocotyl were transferred to soil, and further characterized using the PCR based analysis with 35S promoter specific forward primer and TaSnRK1α gene specific reverse primers. Same selection approach was used in T1 and T2 generation and grown till maturity. PCR based positive plants at T3 homozygous lines were further used for functional characterization. Protein abundance of TaSnRK1α in Arabidopsis overexpression lines was checked using the wheat anti-TaSnRK1α antibody in comparison to the wild type Col-0.

### Arabidopsis mutant complementation using wheat SnRK1

To characterize the function of TaSnRK1α in Arabidopsis, a T-DNA insertion mutant of the catalytic unit of SnRK1 (*KIN10*) in Arabidopsis was obtained from ABRC (Ohio State University, USA). For complementation in *Arabidopsis thaliana (kin10)*, full length CDS *TaSnRK1*α was cloned in plant expression vector pCAMBIA 1302 using the restriction enzyme sites NcoI on both ends using the Gibson assembly (NEB, USA) and empty vector pCAMBIA 1302 was taken as control. Primer details are provided in Supplementary Table 1. Plasmid 35S:TaSnRK1α and empty vector were transformed into *Agrobacterium tumefaciens* (GV1301) cells. For Arabidopsis transformation, seeds of mutant *kin10* and Col-0 were grown in a plant growth chamber at 22 °C, with 16 h light and 8 h dark period till the flowering stage. Plants were transformed by the floral dip method (Zhang *et al*., 2006) and seeds were collected after maturation. Multiple transformants were selected on half MS media with 30 mg/ml hygromycin selection. The seedlings with long hypocotyl and green in colour were selected and transferred to the soil after 4 weeks of selection. T1 and T2 plants were selected similarly and grown till maturity. Transgenic plants were selected using the PCR-based genotyping approach using 35S promoter specific forward primer and TaSnRK1α gene specific reverse primer. The PCR-based positive plants at T3 homozygous generation were further used for functional characterization. An immunoblot assay was performed to check the transgenic Arabidopsis lines. Protein from each line was isolated using the PBS buffer and immunoblotted with the synthesized anti-TaSnRK1α antibody, with the help of a wet-transfer buffer system.

### Starch estimation from Arabidopsis rosette

For iodine staining in Arabidopsis, 4 weeks old rosette for each genotype was collected at the end of day and chlorophyll was destained using 80 % ethanol. Excess ethanol was removed by water prior to iodine staining in Lugol’s solution (KI/I2 solution; Sigma-Aldrich). Extra Lugol staining was removed by water and rosettes were imaged.

For total starch estimation using enzymatic reaction, starch was extracted using the method described by Smith and Zeeman, (2006) from *TaSnRK1*α Arabidopsis O/E lines, *kin10*+*TaSnRK1*α complemented leaves along with Col-0, and mutant *kin10*. Briefly, 4 weeks old rosette was harvested at the end of the light period and grounded into powder using 0.7 M perchloric acid. The extracted suspension was pelleted by centrifugation at 4 °C and washed three times with 80 % ethanol and suspended in distilled water. The starch was gelatinized by boiling at 95 °C for 15 min and then digested using α-amylase and amyloglucosidase (Megagyme) at 37 °C for 6-7 hours. The starch content (glucose equivalent) was estimated using the DNSA method (Rajbhar *et al*., 2015). The total amount of glucose was calculated by subtracting the digested sample from the undigested sample and the liberated glucose concentration was estimated using the glucose standard.

### Quantitative real-time expression analysis

Rosette for the Col-0, *kin10* mutant, *TaSnRK1*α O/E lines and *kin10*+*TaSnRK1*α *c*omplemented were collected at the end of the light period. Total RNA was extracted manually using the TRIzol® Reagent (Invitrogen™). RNA integrity was checked on 1.5% agarose gel and further used for cDNA preparation. A total of 2 µg of total RNA was used for cDNA preparation using the Invitrogen SuperScript III First-Strand Synthesis System SuperMix (Thermo Fisher Scientific) according to manufactured protocol. The sequence of Arabidopsis starch synthesis genes were retrieved from the TAIR database, and gene specific primers were synthesized using Primer3 (Untergasser *et al*., 2012). Primer details are provided in Supplementary Table 1. The qRT-PCR was performed using TB Green® Premix Ex Taq (Takara, Tokyo, Japan) in three biological replicates on a CFX96 thermocycler (BIO-RAD, USA) using the method described previously (Madhawan *et al*., 2020; Kumar *et al*., 2022). The Arabidopsis actin was used as an internal control for the normalization of expression. The relative qRT-PCR expression was calculated by the method described by Livak and Schmittgen (2001), and fold change values were converted into log_2_ fold change scale.

### Starch granule isolation from Arabidopsis leaves

To see the structural differences in the starch granule morphology, starch granules were isolated from the 4-weeks old Arabidopsis plants according to the previously described protocol by Seung *et al*., (2017). Briefly, 4 weeks old plants were collected at the end of the day and ground into fine powder using liquid nitrogen. The powder was dissolved into the extraction buffer (200 mM HEPES-NaOH, pH- 7.4, 0.4 mM EDTA and 0.05% Triton X- 100). The dissolved product was filtrated using the 22 µm filter and the filtrate was centrifuged at 4 °C at 1500 g for 5 min. After centrifugation, the supernatant was discarded and the pellet was dissolved into 40 ml extraction buffer. The dissolved product was filtered using the 100 µM and 30 µM mesh filter and the filtrate was centrifuged at the same conditions. After centrifugation, sediment was dissolved into the 2 ml extraction buffer and slowly pipetted onto Percoll® for density-based separation. This filtrate was centrifuged at 4 °C at 1500 g for 15 mins. The sedimented product containing starch was washed 3 times with chilled water and after removing the excess water, dried using a lyophilizer.

Morphological and structural variations in starch granules were investigated by field emission scanning electron microscopy (FE-SEM). Starch samples were coated with 10 nm gold particles with 15 ma current for 1 min using Quorum Q150T and imaged using Apero S (Thermo Scientific, USA). To characterize the starch extracted from Arabidopsis rosette, FTIR spectra was performed using the infrared spectrometer 536 (ATR-FTIR) (PerkinElmer UATR Spectrum Two). Spectra of starch were recorded between 2000 to 500 cm^-1^. Spectra for the samples previously normalized by noise and baseline correction.

### Enzymatic activity assay of AGPase in Arabidopsis

For measuring AGPase activity, 0.3 g rosette sample from each Arabidopsis genotype was ground into powder in extraction buffer (100 mM HEPES, pH-7.5, 8 mM MgCl_2_, 2 mM EDTA, 1 mM DTT and 12.5 % glycerol) at 4 °C and centrifuged at high speed. The fresh supernatant was used for activity assay. The AGPase activity was measured in accordance with the method of Kulichikhin *et al*., (2016) using ADP-glucose and inorganic pyrophosphate as a substrate and recording the final end product at absorbance (A_340_).

### In-Gel enzyme activity assay of AGPase

To see the enzymatic activity of AGPase in Arabidopsis plant leaves, Zymogram was performed as previously described by Huang *et al*., (2014). Briefly, 3 gm leaves of 4-weeks old plants from wild type (Col-0), *kin10* mutant, *TaSnRK1*α O/E lines, and *kin10*+*TaSnRK1*α Arabidopsis rosettes were collected and grounded in the zymogram extraction buffer using the pre-chilled mortar-pestle on ice. The buffer contained 100 mM Tris-HCl of pH 7, 40 mM β-mercaptoethanol, 100 mM KCl, 10 mM MgCl_2,_ and 15% glycerol. After being ground in extraction buffer, the homogenate was centrifuged at 14000 RPM for 45 min at 4 °C and the supernatant was proceeded with enzymatic activity. For enzymatic activity, equal amount of protein was used and protein concentration was determined using the BCA Protein Assay Kit (Puregene, New Delhi, India). A total of 80 µg protein from each genotype was used further and loaded on the native-acrylamide gel and run in Laemmli buffer without SDS at 4 °C at 90 V for 4-5 hours. After running, the gel was incubated in dark at 37 °C for overnight in incubation mixture, containing 100 mM Tris-HCl, pH 8, 5 mM β-mercaptoethanol, 10 mM MgCl_2_, 5 mM CaCl_2_, 5 mM Glucose-1-phosphate, 5 mM ATP and 10 mM PGA. For the control reaction, the incubation mixture omitted the Glucose-1-phosphate. After incubating overnight, white PPi precipitated bands were visualized and photographed on black background.

### Yeast two-hybrid assay

For the yeast-two hybrid assay, full length CDS of *TaSnRK1*α and AGPase large subunit (LS) was cloned into pGBKT7 and pGADT7 respectively, and transformed into Y2H Gold strain by Li-Ac method. Primer details are provided in Supplementary Table 1. Positive transformants were then checked for self-autoactivation. For autoactivation test, growth of TaSnRK1α cloned in pGBKT7 vector was checked in double dropout media (DDO; lacking/-trp/-leu) and triple dropout media (TDO; lacking/-trp/-leu/-his) media with X-alpha-gal. For yeast one-on-one interaction, mixed recombinant plasmids were transformed together and plated on DDO media. To see the positive interaction, transformants were selected on DDO media, and quadruple dropout media (QDO; lacking/-trp/-lue/-His/-Ade) supplemented with Aureobasidin and X-alpha-Gal.

### Bimolecular Fluorescence Complementation (BiFC)

To identify the protein-protein interaction between wheat AGPase large subunit (LS) and wheat TaSnRK1α, a BiFC assay was performed using the split-YFP system in tobacco leaves. For BiFC assay, CDS of *TaSnRK1*α was cloned into pUC-SYNE with the help of restriction enzymes sites BamHI and XhoI, while wheat AGPase LS was cloned into pUC-SYCE with the help of Gibson assembly master mix (NEB), using BamHI as restriction enzyme (Sharma *et al*., 2022). Primer details are provided in Supplementary Table 1. Both TaSnRK1α and AGPase LS constructs and empty vectors were transformed further into Agrobacterium cells GV 1301 strain. The constructs were co-infiltrated in tobacco leaves and YFP signals were assessed using the laser scanning confocal microscopy on Carl Zeiss confocal microscope LSM880 (Germany).

### Statistical analysis

The number of biological replicates and number of samples with statistical details for each experiment are described in the corresponding figure legends. All the statistical analysis and graphs were made using Graphpad-Prism version 8 (GraphPad Software, Inc., CA, US) and Microsoft Excel 2016.

## Results

### In silico analysis of TaSnRK1α in wheat

In our previous study (Kumar *et al*., 2022), we identified TaSnRK1α regulating starch biosynthesis in wheat. In silico analysis identified TaSnRK1α belongs to plant non-specific Ser/Thr kinase (SnRK1) and is ortholog of yeast SNF-1 and animal AMPK. TaSnRK1α is encoded by the (TraesCS3A02G282800, TraesCS3B02G316500, and TraesCS3D02G339800), encoding 349 amino acids protein sequence and there was no difference in amino acid sequence of three homeologues. Domain analysis using SMART database and InterProScan predicted three functional domains in TaSnRK1α, namely, ATP-binding domain (20-43 a.a.), Ser/Thr kinase domain (133-145 a.a.), and UBA domain (288- 328 a.a.) (Fig. 1A). The presence of UBA domain with Ser/Thr kinase domain is the characteristic feature of SnRK1.

**Fig. 1:**
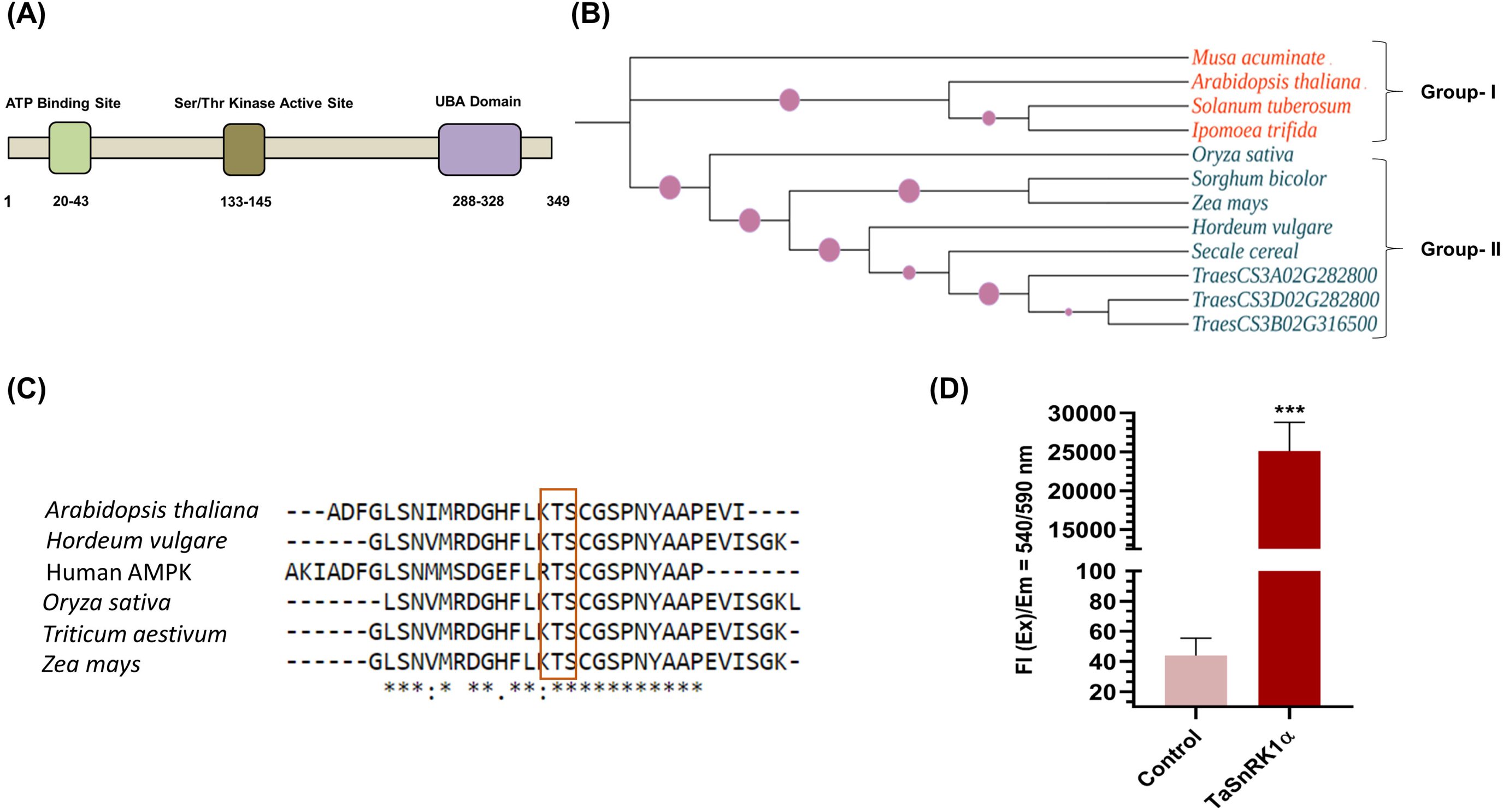
In silico analysis and protein kinase assay of TaSnRK1α. **(A)** Domain architecture of TaSnRK1α, showing three different domains in different colors. Numeric values show the position of amino acids respective to each domain. **(B)** Phylogenetic analysis of TaSnRK1α with other plant species. Monocots clustered together in group I whereas dicots clustered together in group II. **(C)** Amino acid sequence alignment of SnRK1 in different plant species. The box shows the conserved Threonine (Thr) residue present in the sequences. **(D)** The enzymatic kinase activity of TaSnRK1α and control protein (GBSSI) was detected using Universal Fluorometric Kinase Assay at 37 °C for 30 mins. Data are presented as Means ± SEM (n = 3). (T-test; ****, p < 0.0001; n=3). FI: Fluorescence intensity.

Further to see the evolutionary relationship with different plant groups, we performed the phylogeny analysis with the different monocots and dicots family that are rich in starch. In evolutionary study, all the three wheat homeologue grouped together and showed closest similarity with *Secale cereale* and *Hordeum vulgare*. All the dicots fell into group-I and all other monocots fell into group-II (Fig. 1B).

AMPK, SNF-1 and SnRK1 gets activated by T-loop phosphorylation at Thr residues by upstream kinases (Crozet *et al*., 2014). Multiple sequence alignment of TaSnRK1α, protein sequences from animal AMPK and various other plant specific SnRK1 showed the presence of T-loop in TaSnRK1α. In AMPK, T-loop was present at Thr172 while in cereals (including wheat, rice, maize and barley) T-loop was present at Thr170 position, whereas in case of Arabidopsis T-loop was present at Thr175. Thr loop consist of quartet amino acids (LKTS; letter denotes amino acids) while in human it consists of LRTS (Fig. 1C). All the results indicate the TaSnRK1α belongs to plant SnRK1.

### TaSnRK1α showed kinase activity in vitro

To study the kinase properties of the TaSnRK1α, in vitro kinase assay was performed using the recombinant protein expressed in *E. coli*. AMARA peptide was previously identified as the universal substrate of the various kinases. So, in the current study we used AMARA peptide as the TaSnRK1α substrate and analysed the enzymatic activity. Enzymatic activity was measured as amount of ADP released through the reaction. Amount of ADP released was measured through Universal Fluorometric Kinase Assay Kit (Sigma). In house recombinant protein GBSSI was used as control protein in the assay. Fluorescence intensities in the reaction indicated the kinase activity of TaSnRK1α. High kinase activity was observed in TaSnRK1α while GBSSI did not show any kinase activity in the reaction (Fig. 1D).

### *TaSnRK1*α functionally complement yeast Δ*snf1*

To investigate the function of wheat *TaSnRK1*α, yeast complementation assay was performed in yeast mutant Δ*snf1*. SNF1 is the ortholog of SnRK1 in yeast and its mutant Δ*snf1* is deficient to utilize the sucrose as a carbon source and glycogen accumulation. To characterize the function, full length CDS of *TaSnRK1*α was expressed in the yeast wild strain CEN.PK and mutant Δ*snf1* and growth was observed. We found that the cells expressing Δ*snf1+TaSnRK1*α were able to grow on sucrose media while Δ*snf1* cells were able to utilize only glucose media and showed growth defect on sucrose media after growing plates on appropriate conditions (Fig. 2A). Wild type strain CEN.PK and CEN.PK+*TaSnRK1*α showed normal growth phenotype on both glucose and sucrose media. Therefore, the TaSnRK1α was able to rescue growth defect of Δ*snf1*.

**Fig. 2:**
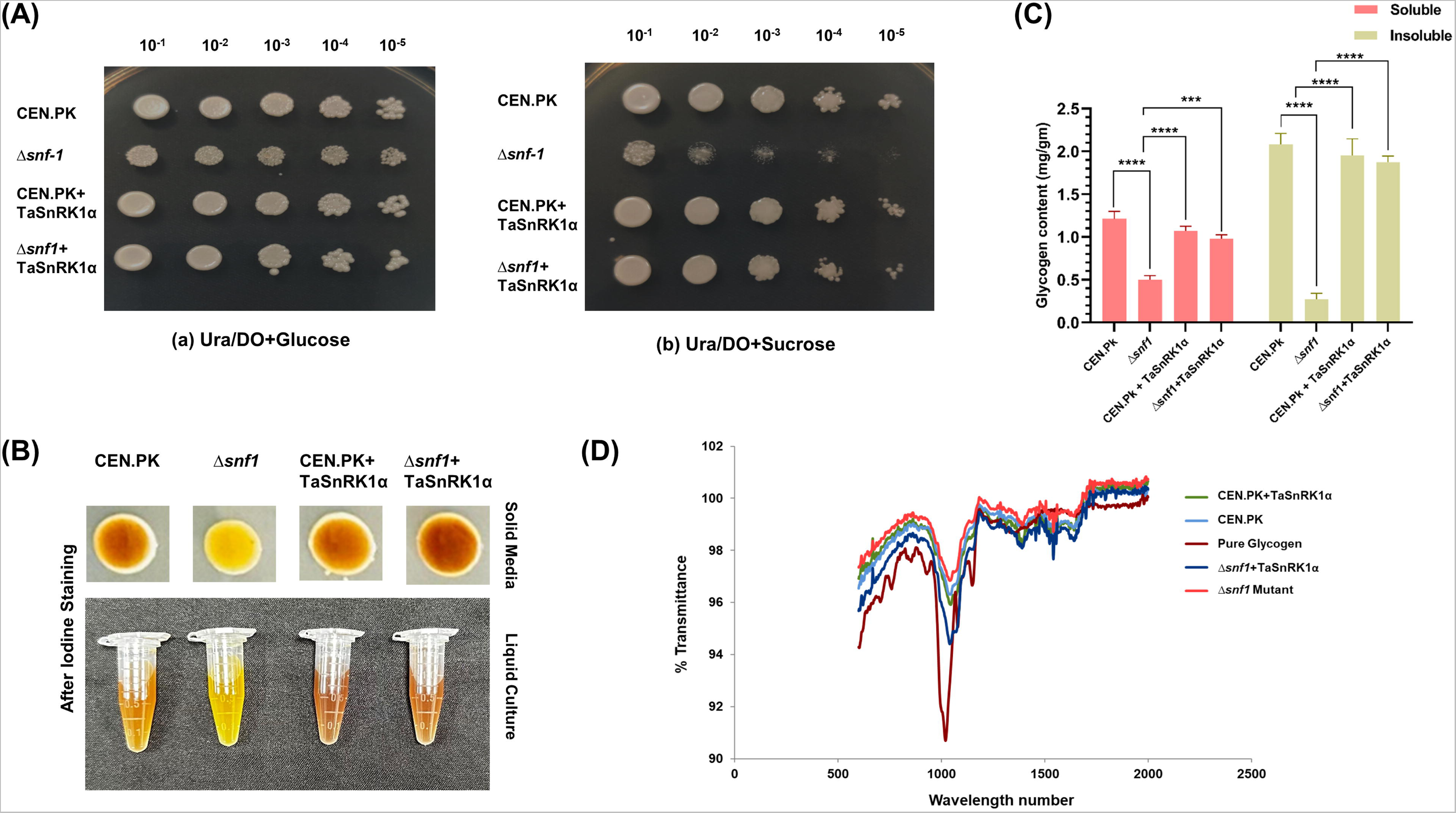
TaSnRK1α complements growth defect and promotes glycogen accumulation in yeast Δ*snf1* mutant. **(A)** Spotting assay of yeast strain CEN.PK and Δ*snf1* mutant transformed with TaSnRK1α. Serially diluted cultures of each transformant were spotted on minimal media lacking Uracil in the presence of either glucose or sucrose.10 µl of each dilution were spotted on plates and allowed to grow at 30 °C for 3 days. Ura/DO; Uracil dropout media **(B)** Quantification of soluble and insoluble glucans extracted from yeast transformant grown for 2 days in liquid cultures. Glycogen content was estimated in mg/ml fresh weight. Soluble and insoluble fractions are shown in different colors. (Two-way ANOVA, with Tukey-test multiple comparison test; ***, p < 0.001, **** p < 0.0001; n=3). **(C)** Iodine staining of each yeast transformant grown on SC-gal plates and liquid growth medium with 2% galactose and 1% raffinose. The intensity of the color represents the glycogen content. **(D)** FTIR spectra of yeast glycogen extracted from each transformant along with commercially available Oyster glycogen, scanned from 500 to 2000 cm-^1^. Corresponding peak at 1000 cm-^1^ shows the presence of a glycosidic bond of glycogen in each sample.

Further, to assess role of TaSnRK1α in glucan accumulation in yeast cells, glycogen content was estimated qualitatively and quantitatively. We assessed the glycogen content in yeast cells using histochemical studies. Lugol solution stains glycogen brown in color in yeast cells and color intensity is directly proportional to the glycogen content. After staining, cells from Δ*snf1* showed less brown and yellow color because of less glucan production in the cells while wild type (CEN.PK), CEN.PK+*TaSnRK1*α and complemented Δ*snf1*+*TaSnRK1*α show more brown color cells (Fig. 2B). Similar results were found when cells from liquid culture were observed under light microscopy. Liquid culture cells from Δ*snf1* stained less while, CEN.PK, CEN.PK+*TaSnRK1*α and complemented Δ*snf1*+*TaSnRK1*α stained browner in color (Supplementary Fig. 1).

To further validate the histochemical results, glycogen was quantified using DNSA method. Three different colonies from each of the transformants were used for glycogen assay. It was observed that glycogen content of mutant Δ*snf1* was lower than CEN.PK. Whereas in Δ*snf1*+*TaSnRK1*α complemented cells, TaSnRK1α promotes glycogen accumulation in yeast cells and glycogen content of complemented cells was comparable to CEN.PK. Insoluble and soluble fraction of mutant Δ*snf1* was significantly lower than the CEN.PK and Δ*snf1+TaSnRK1*α complemented cells. Insoluble fraction of Δ*snf1,* CEN.PK and Δ*snf1+ TaSnRK1*α was found to be 0.27 mg/ml, 2.08 mg/ml and 1.87 mg/ml respectively, meanwhile, the soluble fraction was found to be 0.50 mg/ml, 1.21 mg/ml and 0.98 mg/ml respectively (Fig. 2C).

Moreover, FTIR spectroscopy showed the characteristic spectra of glycogen with insoluble fractions obtained from each of the transformant. Prominent peak of C-O-C glycosidic bond at 1000 cm^-1^ was observed i.e., characteristics peak of glycogen in each yeast transformants (Fig. 2D). Overall, the above results indicate role of TaSnRK1α in carbohydrate metabolism and glycogen accumulation in yeast cells.

### Effect of ATP binding and UBA domain on TaSnRK1α activity in yeast

To study the effect of key domain ATP-binding domain and UBA domain on the function of TaSnRK1α, complementation assay was performed in yeast. For the complementation assay, *TaSnRK1*α lacking one of the domains were over-expressed in Δ*snf1* yeast mutant. The *TaSnRK1*α lacking any of the UBA or ATP binding domain failed to complement the mutant phenotype on sucrose media, on contrary to the wild type (Fig. 3A). So, these results reflect both the domains are required for the activity of the TaSnRK1α.

**Fig. 3:**
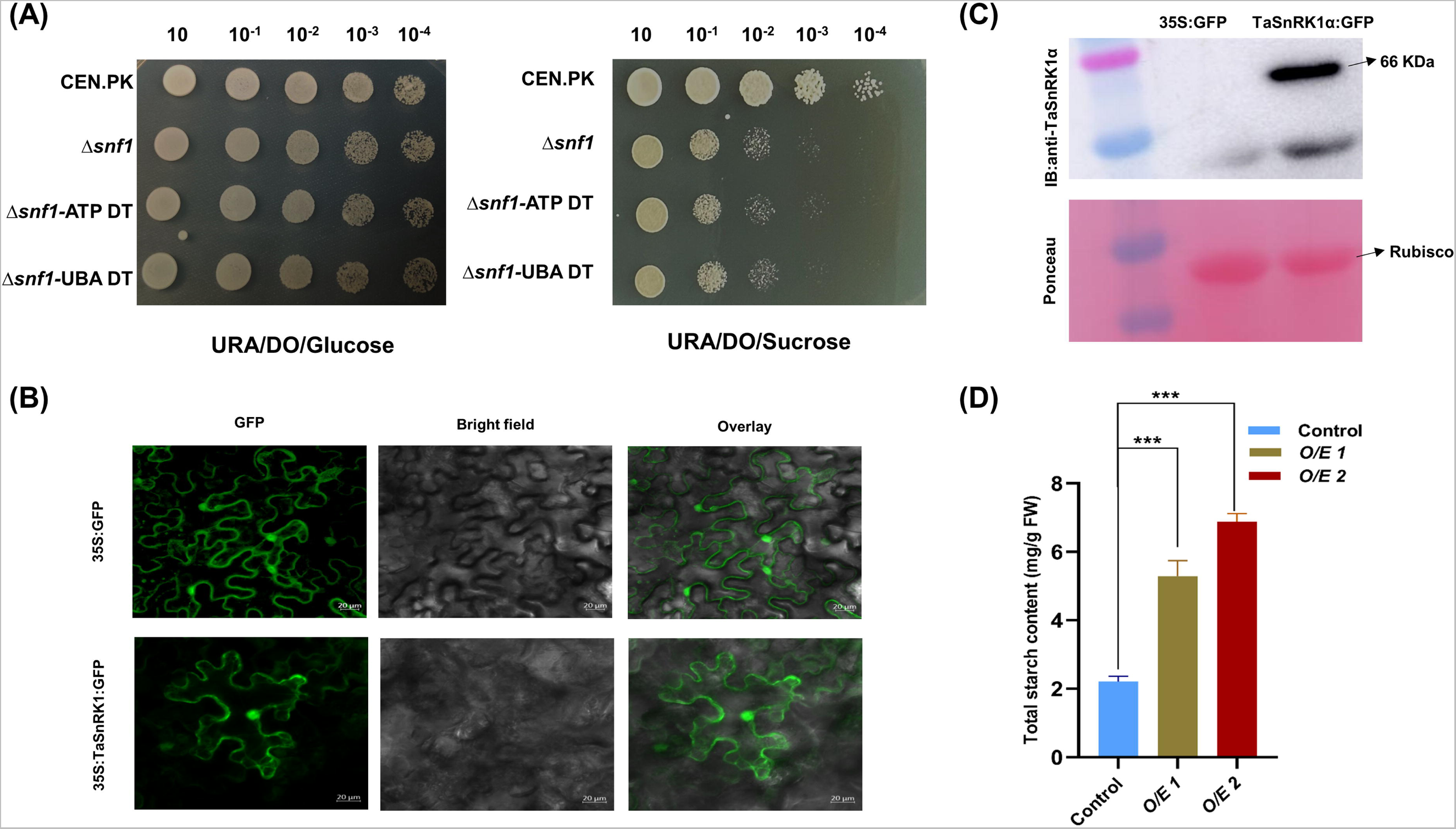
Domain deletion analysis of TaSnRK1α in yeast system, sub-cellular localization, and transient starch estimation in tobacco. **(A)** TaSnRK1 α truncated sequence lacking either ATP-binding domain or UBA domain were generated by PCR mediated deletion approach and transformed into pYES2. Spotting assay was performed on minimal media lacking uracil in presence of either glucose or sucrose. Growth of serially diluted cultures was assessed after incubating at 30 °C for 3 days. Ura/DO; Uracil dropout media **(B)** Sub-cellular localization of 35S:TaSnRK1α:GFP recombinant protein was assessed in tobacco leaves. Agrobacterium transformed 35S:TaSnRK1α:GFP was infiltrated into the 4-week-old tobacco leaves, and GFP signal was observed under confocal microscopy after 2 days of infiltration. An empty vector was used as a positive control. All the GFP images were taken with a gain % of 700 to maintain consistency. **(C)** Immunoblot showing TaSnRK1α:GFP fused protein abundance in the 35S:TaSnRK1α:GFP infiltrated leaves and control leaf with empty vector. Wheat anti-TaSnRK1α specific antibody was used for immunoblot assay in 1:5000 dilution and ponceaus S Staining shows the equal intensity Rubisco in the western blot. **(D)** Total starch content in transiently expressed TaSnRK1α in *N. benthamiana* leaves. Total starch was quantified using enzymatic digestion from two independent leaves (O/E 1 and O/E 2). An empty vector infiltrated leaf was used as an internal control. Error bars denote standard deviation. (Ordinary One-way ANOVA; ***, p < 0.001; n=3). O/E: TaSnRK1α over-expression.

### Sub-cellular localization of TaSnRK1α in tobacco leaves

To understand the site of function of the TaSnRK1α, we analysed the sub-cellular localization of TaSnRK1α in tobacco leaves. The TaSnRK1α was predicted to localize in nucleus, cytoplasm and mitochondria using the BUSCA 3 web-server. Upon transient expression, the 35S:TaSnRK1α:GFP recombinant protein showed the GFP signal in nucleus and cytoplasm, while the control signal (CaMV:35S:GFP) showed GFP signal all over the cells (Fig. 3B).

To detect the protein abundance of TaSnRK1α fused protein with GFP at C-terminal, immunoblot assay was performed using wheat specific anti-TaSnRK1 antibody. In immunoblot assay, protein abundance of TaSnRK1α:GFP fused protein was detected and showed the band of 66 KDa (Fig. 3C).

### Transient overexpression of *TaSnRK1*α accumulate starch in tobacco leaves

To investigate the function of TaSnRK1α in starch biosynthesis in planta, CDS of *TaSnRK1*α was transiently expressed in tobacco leaves under constitutive promoter 35S. Total starch content was quantified in Agrobacterium infiltrated leaves of 35S:*TaSnRK1*α along with control leaf infiltrated with empty pCAMBIA1302 vector. After 7-days of infiltration, starch content was measured from infiltrated leaves from two independent plants using DNSA method and observed high starch content in *TaSnRK1*α transiently expressed leaves in comparison to the control leaves. Total starch content of the control leaf was 2.21 mg/g fresh weight (FW) while for transiently expressed leaves it was 5.29 mg/g FW and 6.88 mg/g of FW from two different plant leaves. Starch content of the *TaSnRK1*α transient expressed leaves were 2.3 and 3.1 fold higher than the control leaves and therefore, depict the role of TaSnRK1α in starch biosynthesis in vivo (Fig. 3D).

### *TaSnRK1*α heterologous overexpression in Arabidopsis accumulate high starch

To reveal the function of TaSnRK1α, CDS of *TaSnRK1*α was overexpressed in Arabidopsis ecotype Col-0. The overexpressed homozygous T3 lines were prepared and lines were confirmed through DNA amplification. Gene specific primers showed TaSnRK1α specific band in agarose gel that was not present in the wild type plants (Supplementary Fig. 2). To confirm the TaSnRK1α transgenic lines, immunoblot assay was also performed using wheat specific anti-TaSnRK1α antibody and band of TaSnRK1α corresponding to 39 KDa was observed in transgenic Arabidopsis proteins (Fig. 4A).

**Fig. 4:**
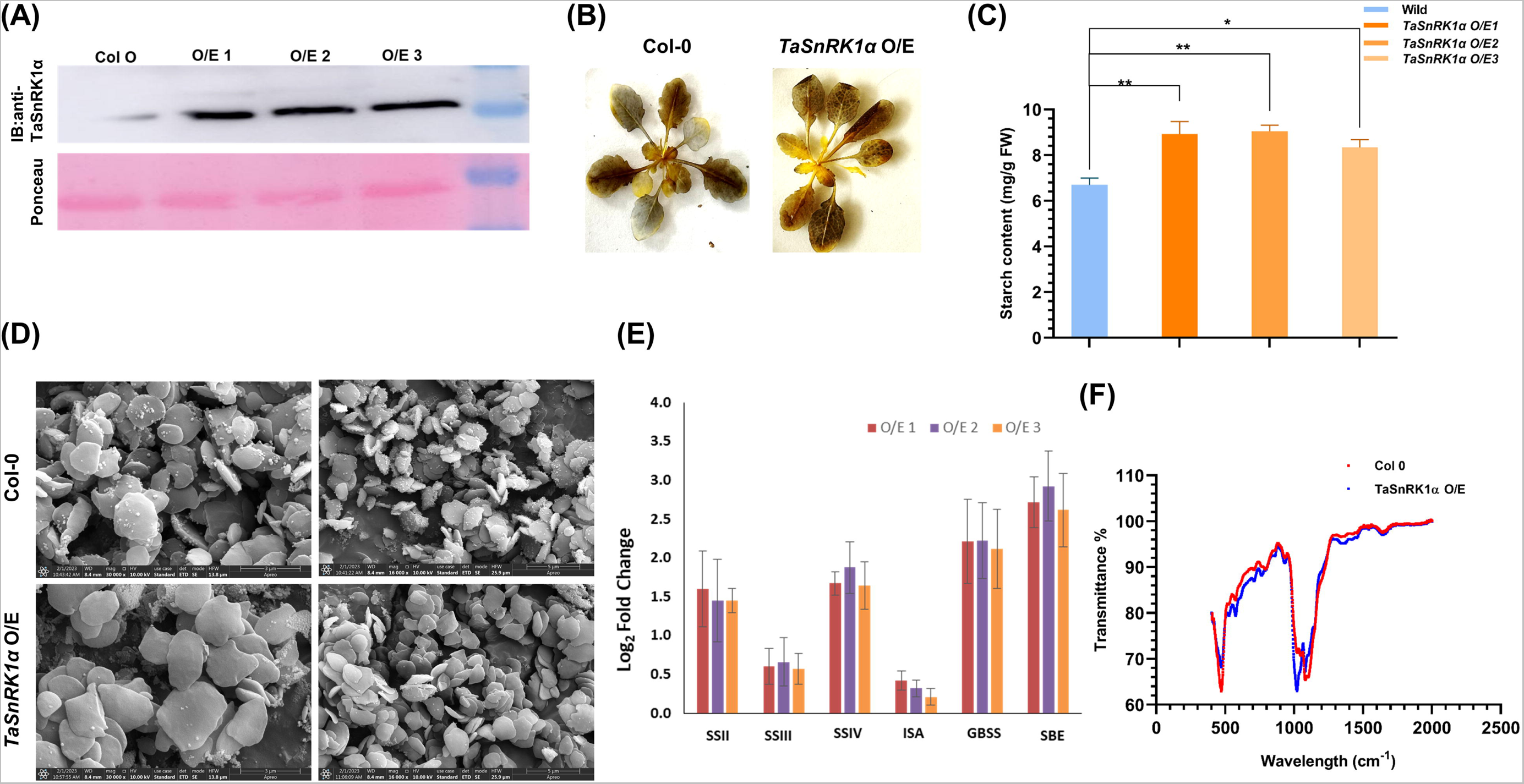
TaSnRK1α overexpression accumulates starch in transgenic Arabidopsis. **(A)** Immunoblot analysis showing protein abundance of TaSnRK1α in Arabidopsis overexpression (O/E) transgenic lines. Three independent O/E lines designated as O/E 1, O/E 2 and O/E 3 along with Col-0 as control were used. Immunoblot analysis was performed using the anti-TaSnRK1α antibody in 1:5000 dilution with a wet transfer system and ponceaus S Staining shows the equal intensity Rubisco in the western blot. **(B)** Iodine staining showing difference in starch content in rosette of Col-0 and TaSnRK1α O/E line. **(C)** Starch content estimated in TaSnRK1α O/E transgenic lines and wild type Col-0. Three independent lines (TaSnRK1α O/E 1, TaSnRK1α O/E 2 and TaSnRK1α O/E 3) were used for starch estimation and the experiment was repeated three times. (Two-way ANOVA, with Dunnett’s multiple comparison test; *, p < 0.05, **, p < 0.01). **(D)** Comparison in the starch granule morphology in Arabidopsis TaSnRK1α O/E line and Col-0. Starch granules were purified to observe under scanning electron microscopy and images were captured at 16000x and 30000x magnification. **(E)** qRT expression of starch pathway genes in TaSnRK1α O/E lines in comparison to wild type Col 0. All the data are represented as mean[±[SD from three biological and three technical replicates.**(F)** FTIR spectra of starch granules isolated from TaSnRK1α O/E line and wild type Col-0 showing a characteristic peak at 1000 cm-^1^ corresponding to the presence of a glycosidic bond in each sample.

To assess the role of TaSnRK1α in starch biosynthesis pathway, starch accumulation was qualitatively and quantitatively estimated in transgenic O/E and wild type Col-0. Upon iodine staining, rosette of O/E line show more iodine uptake and stained browner in color in comparison to the Col-0 (Fig. 4B). The total starch content of wild type was 7.03 mg/g FW where in transgenic O/E lines it was significantly high i.e., 8.92, 9.38 and 8.67 mg/g of FW (Fig. 4C).

To see the structural variation in starch granules of *TaSnRK1*α O/E lines and Col-0, FE-SEM was performed. Under the SEM analysis, starch granules of the overexpression lines were relatively larger in size in comparison to the Col-0 (Fig. 4D). To explore the potential regulatory effect of TaSnRK1α, expression of starch pathway genes was assessed in the O/E lines in comparison to the wild type plants. In qRT-PCR, GBSSI and SBE showed higher expression in the O/E lines, starch synthases showed moderate up-regulation in comparison to the wild type Col-0 where no significant difference was found in the expression of ISA (Fig. 4E).

To characterise the starch granules, FTIR analysis was performed and significant peak of glycosidic bond present in starch was observed at 1000 cm^-1^. Moreover, peak intensity of *TaSnRK1*α O/E was higher in comparison to the Col-0, which reflects presence of more glycosidic bonds in O/E lines corresponding to the more starch content (Fig. 4F). These results suggest that *TaSnRK1*α overexpression may contribute to high starch accumulation in Arabidopsis.

### *TaSnRK1*α restored starch accumulation in *kin10* Arabidopsis mutant

KIN10, a catalytic subunit of SnRK1 complex in Arabidopsis, shows similarity with TaSnRK1α. In order to establish the function of TaSnRK1α, Arabidopsis *kin10* mutant was complemented with *TaSnRK1*α. Stable transformed lines expressing wheat *TaSnRK1*α under the constitutive promoter CaMV:35S were generated and all the analysis was performed at T3 generation. Initially, seeds of the *kin10* mutants were confirmed by checking the expression of KIN10 and the qRT-PCR results show no expression of KIN10 in *kin10* mutant in comparison to Col-0 (Fig. 5A).

**Fig. 5:**
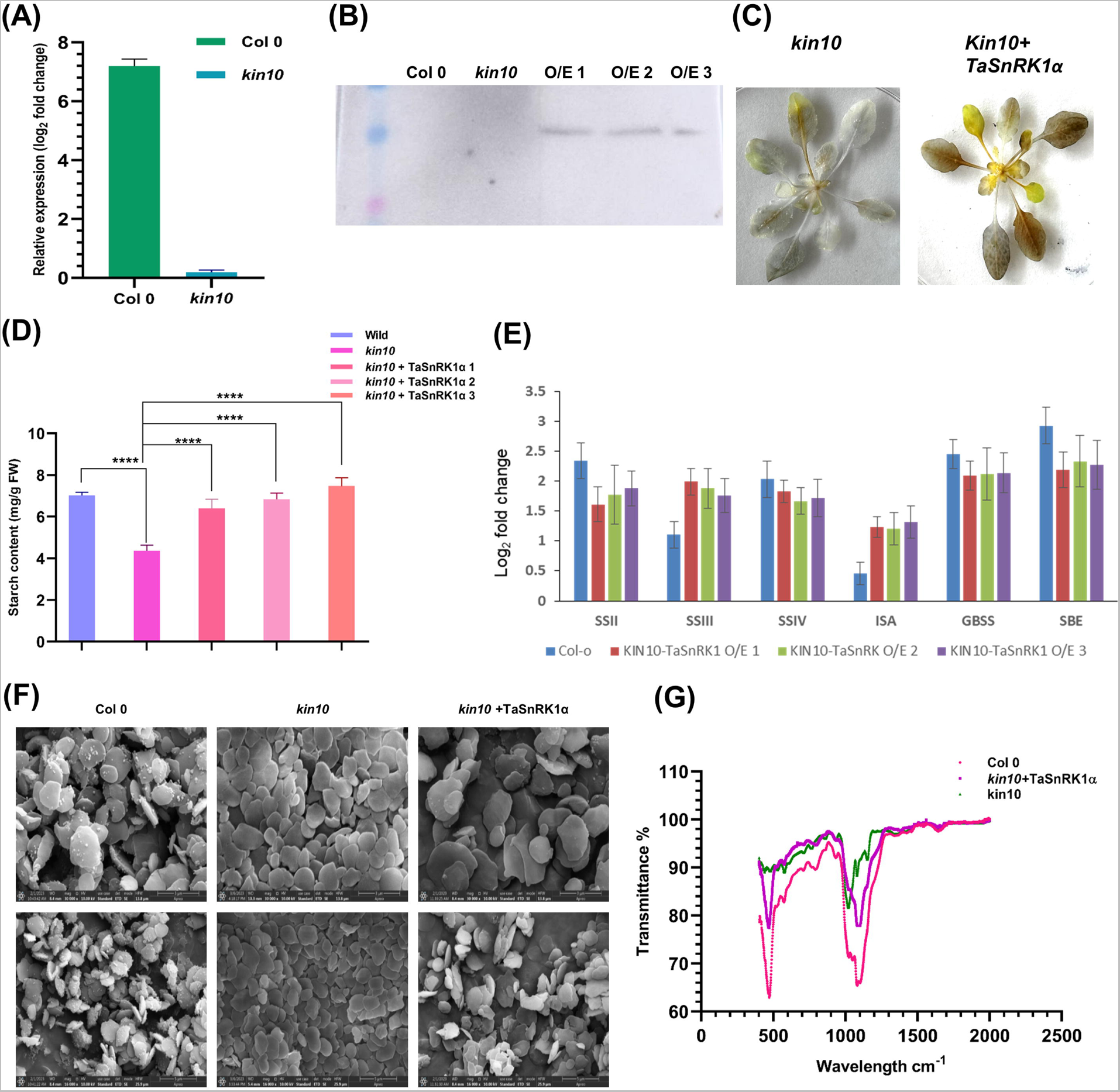
TaSnRK1α complement *kin10* mutant in Arabidopsis and accumulate starch. **(A)** qRT expression of KIN10 in Col-0 and mutant *kin10*. Data are represented as mean[±[SD from three biological and three technical replicates **(B)** Immunoblot analysis showing protein abundance of TaSnRK1α in wild type Col-0, mutant *kin10,* and *kin10*+TaSnRK1α complemented transgenic lines. Immunoblot assay was performed using wheat anti- TaSnRK1α antibody in 1:5000 dilution with a wet transfer system with a wet transfer buffer system. **(C)** Iodine staining shows the difference in starch content in a rosette of Col-0 and *kin10*+TaSnRK1α complemented line. **(D)** Starch content estimation in wild-type Col-0, mutant *kin10,* and *kin10*+TaSnRK1α complemented transgenic lines. Three independent lines (*kin10*+TaSnRK1α 1-3) were used for starch estimation and the experiment was repeated three times. (Two-way ANOVA, with Dunnett’s multiple comparison test; *, p < 0.05, **, p < 0.01) **(E)** qRT expression of starch pathway genes in *kin10*+TaSnRK1α complemented and Col-0 in comparison to mutant *kin10*. All the data are represented as mean[±[SD from three biological and three technical replicates. **(F)** Comparison in the starch granule morphology in Arabidopsis *kin10* mutant with Col-0 and *kin10*+TaSnRK1α line. Starch granules were purified to observe under scanning electron microscopy and images were captured at 16000x and 30000x magnification. **(G)** FTIR spectra of starch granules isolated from *kin10*, *kin10*+TaSnRK1α line and wild type Col-0 showing a characteristic peak at 1000 cm-^1^ corresponding to the presence of a glycosidic bond in each sample.

*kin10+TaSnRK1*α complemented transgenic lines were confirmed using PCR-based genotyping approach and TaSnRK1α specific band was observed in complemented lines while band was not present in mutant *kin10* (Supplementary Fig. 3). Further, *kin10+TaSnRK1*α transgenic lines were confirmed with the immunoblot assay using wheat specific anti-TaSnRK1α antibody and result show TaSnRK1α protein specific band at 39 KDa in transgenic lines while no bands were observed in wild type Col-0 and mutant *kin10* (Fig. 5B). To gain the insight of TaSnRk1α in starch biosynthesis using qualitative method, iodine staining was performed. Upon staining with Lugol’s solution mutant *kin10* show very less iodine-stained rosette while *kin10*+*TaSnRK1*α show more brown color (Fig. 5C). Starch accumulation was checked in *kin10*+*TaSnRK1*α transgenic lines along with Col-0 and mutant *kin10*. Upon estimation mutant *kin10* showed less starch (4.3 mg/g FW) in comparison to the Col-0 (7.1 mg/g FW) and *kin10*+*TaSnRK1*α complemented lines, whereas starch content of three independent *kin10*+*TaSnRK1*α complemented lines (6.52, 6.85 and 7.55 mg/g FW) were comparable to Col-0 (Fig. 5D). Upon qRT-PCR analysis, expression of key genes like GBSSI, SBE and starch synthases were found to be significantly up-regulated whereas ISA showed no significant difference in *kin10*+*TaSnRK1*α in comparison to the *kin10* (Fig. 5E).

Starch granules variation in morphology was checked using FE-SEM. *kin10* showed smaller starch granules in comparison to *kin10*+*TaSnRK1*α and Col-0. While *kin10*+*TaSnRK1*α showed similar size of starch granules and were comparable to Col-0 (Fig. 5F). To characterise the starch isolated from Arabidopsis leaves, FTIR analysis was performed, and starches from all genotypes showed significant glycosidic bond peak at 1000 cm^-1^. Peak intensity of mutant *kin10* was lowest in the spectra followed by *kin10*+*TaSnRK1*α and Col-0, thus reflects less starch content in the mutant lines (Fig. 5G). The overall results suggest the TaSnRK1α ability to complement the function of *kin10* and play role in starch biosynthesis in Arabidopsis.

### Enzymatic activity of the AGPase is modulated in Arabidopsis

The overexpression of SnRK1 was previously shown to be positively correlated with enhanced AGPase activity (Jiang *et al*., 2013; Wang *et al*., 2017; Ren *et al*., 2019). To test if same was true in our TaSnRK1α O/E lines, we determined the AGPase enzymatic activity, AGPase activity was significantly lower in the *kin10* mutant (13 nmol min^-1^ mg^-1^ protein) in comparison to the other genotypes. Activity of AGPase was significantly increased in TaSnRK1α O/E lines (25.28 nmol min^-1^ mg^-1^ protein) followed by Col-0 (20 nmol min^-1^ mg^-1^ protein) and *kin10*+TaSnRK1α (18.42 nmol min^-1^ mg^-1^ protein) (Fig. 6A).

**Fig. 6:**
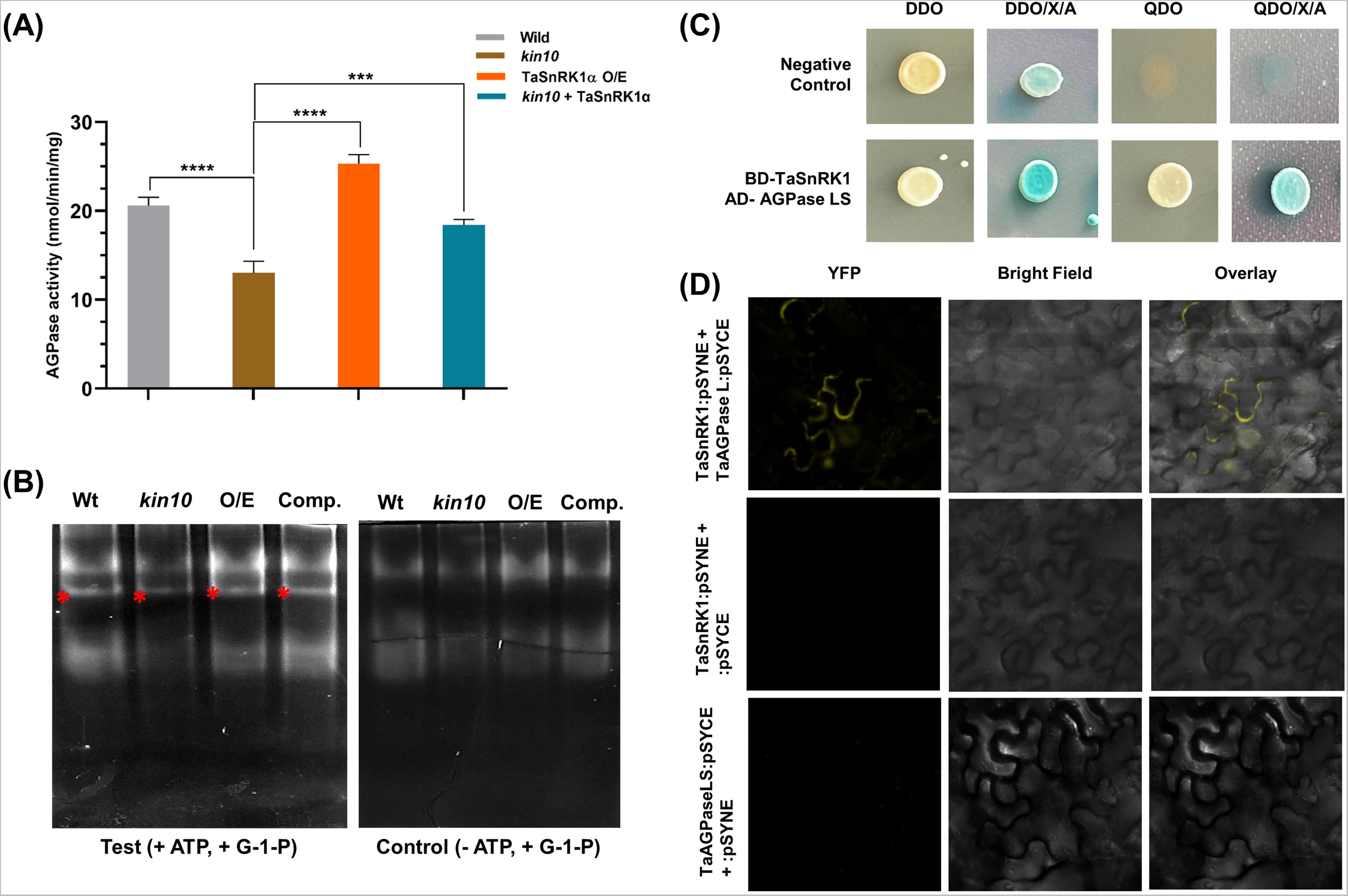
TaSnRK1 interacts with AGPase Large subunit and modulates AGPase activity in Arabidopsis. **(A)** AGPase enzymatic activity was measured in nmol/min/mg of protein extracted from leaf sample of each genotype. Three biological replicates were used for analysis, and the data provided as mean, and bars represents± standard deviation. (Two-way ANOVA with Dunnett’s multiple comparison test; ***, p < 0.001, ****, p < 0.0001). **(B)** Zymogram showing AGPase activity. Zymogram gel showing white precipitated bands generated from enzymatic activity of AGPase. A reaction mixture without ATP was used as a control reaction. Asterisks indicate precipitated bands that require ATP and thus indicate AGPase activity. Wt: wild type; O/E: TaSnRK1α over-expression line; comp: *kin10+* TaSnRK1α complemented line. **(C)** Yeast two-hybrid assay shows the interaction between TaSnRK1α and AGPase large sub-unit. Co-transformed yeast cells were grown on the selective medium SD/-Trp/-Leu (DDO), and SD/-Trp/-Leu/X-alpha-gal/Aureobasidin (DDO/X/A) and interactions were confirmed on SD/-Trp/-Leu/-His/-Ade (QDO), and QDO/X/A. pLam-pGADT7 was used as a negative control. **(D)** Bimolecular fluorescence complementation (BiFC) assays showing the interaction between TaSnRK1α and wheat AGPase Large subunit in tobacco. The proteins were transiently co-expressed with recombinant vectors of BiFC (pSYNE and pSYCE) in N. benthamiana epidermal cells and visualized under confocal microscopy. Reconstitution of yellow fluorescence shows positive interaction Empty vectors with individual constructs were used as controls.

To further verify the activity assay results, in-gel enzyme activity assay (zymogram) was performed. Protein from 4-weeks old rosette from each line was used for analysis and run on native-PAGE. When native gel was incubated in the activity buffer containing Glc-1-P, ATP, positive regulator 3-phosphoglyceric acid and divalent cations, visible white bands were observed due to PPi precipitation. Precipitated PPi bands intensity in zymogram corresponds to the AGPase activity. Light visible bands were seen in Arabidopsis mutant *kin10*, while clear white concentrated PPi precipitated bands were seen in wild type Col-0, *TaSnRK1*α O/E lines and *kin10*+*TaSnRK1*α, with the highest intensity in *TaSnRK1*α O/E and comparable intensity in Col-0 and *kin10*+*TaSnRK1*α demonstrating higher AGPase activity whereas no band was seen in control assay that omitted ATP in it (Fig. 6B). Thus, the above results suggested total AGPase activity was increased due to the expression of TaSnRK1α.

### TaSnRK1α interacts with wheat AGPase large subunit

In order to investigate the physical interaction between TaSnRK1α and AGPase, one-on-one yeast-two hybrid assay (Y2H) was performed. Initially, we examined whether TaSnRK1α could activate itself. Complete CDS sequence of *TaSnRK1*α was cloned into the BD vector and autoactivation was tested in DDO (lacking/-trp/-leu) and TDO (lacking/-trp/-leu/-his) with X-alpha-gal. The results showed that corresponding clone gave white and pale-yellow colonies on DDO with X-alpha-gal while unable to grow on TDO with X-alpha-gal, indicating that TaSnRK1α did not have the property of self-activation.

For interaction assay, TaSnRK1α fused with GAL4 BD and wheat AGPase LS fused with GAL4 AD were co-transformed in yeast strain to implement Y2H mating strategy and tested to grow on double dropout (DDO) and quadruple dropout (QDO) in combination with X-alpha-Gal and Aureobasidin. TaSnRK1α and AGPase LS diploid yeast cells produced from mating, grew on QDO and DDO, and gave blue colonies when supplied with X-alpha-Gal in the dropout medium. Whereas negative control appears white colonies in DDO medium when supplied with X-alpha-gal while no growth was observed in QDO medium (Fig. 6C). Therefore, TaSnRK1α physically interacts with AGPase LS.

Further, we applied the bimolecular fluorescence complementation (BiFC) approach to validate the results *in Planta.* In BiFC assay, TaSnRK1α and AGPase LS were expressed as N-terminal and C-terminal YFP proteins in *N. benthamiana*. Strong cytoplasmic fluorescent YFP signal was expressed in leaves when TaSnRK1α was co-expressed with AGPase LS (Fig. 6D). Whereas no YFP signal was observed when empty vector with single protein was expressed in tobacco leaves. The above results clearly reflected that TaSnRK1α and AGPase LS are capable of physical interaction in yeast and plant cells.

## Discussion

Wheat is one of the important cereal crops in the world and provides major portion of calories in human population. Starch being the main stored carbohydrate in cereal crops provides extended energy in the form of balanced diet. Due to its wide availability, natural biodegradable and hydrophilic properties, starch has important beneficial value in various industrial sectors like food, brewery, textile and chemical industry as a thickener, stabilizer, adhesive and gelling agent (Brennan *et al*., 2008). Therefore, extensive research on detailed understanding of starch metabolism in plants, the genes and their regulation associated with starch biosynthetic pathway is unavoidable. The starch biosynthetic pathway in cereal crops has been studied and is well characterized. However, the regulation of key genes involved in starch biosynthesis remains to be explored. In the present study, we characterized a plant specific Ser/Thr kinase, TaSnRK1α and demonstrated its role in regulating the starch biosynthesis in wheat grain. TaSnRK1α was previously identified in our study to be transcriptionally up-regulated in high amylopectin wheat mutant by transcriptome analysis (Kumar *et al*., 2022).

### Wheat TaSnRK1α is an ortholog of SNF1 and restores growth defect of Δ*snf1* in yeast

Protein homology of TaSnRK1α show similarity with yeast SNF1, animal AMPK and catalytic unit of SnRK1 (*KIN10*) of *Arabidopsis thaliana.* Plant SnRK1, yeast SNF1 and animal AMPK are evolutionary conserved and functions in carbohydrate metabolism (Hey *et al*., 2007), indicating probable role of TaSnRK1α in carbohydrate metabolism. Protein sequence and in silico studies identified three major domains of TaSnRK1α; ATP binding domain, Ser/Thr kinase domain and UBA domain. Presence of UBA and ATP binding domain is key feature of SnRK1 family (Broeckx *et al*., 2016). ATP binding and Ser/Thr kinase domain at N-terminal functions as key catalytic unit while UBA domain at C-terminal functions as regulatory unit of SnRK1. Interestingly, we also found a T-loop at Thr170 which is conserved across the cereal crops. Presence of T-loop and its phosphorylation is important for SnRK1 activity (Baena-González *et al*., 2007; Martínez-Barajas *et al*., 2011). However, the position of T-loop may vary in different system such as Thr172 in Arabidopsis and Thr175 in animals. Overall results indicate, the topology of TaSnRK1α was similar to the previous reports.

*TaSnRK1*α successfully complemented the Δ*snf1* mutant phenotype in yeast and rescued the growth defect on sucrose medium, indicating the utilization of sucrose as a carbon source. The complementation may be attributed probably by activation of the downstream genes like; SUC2 gene, responsible for utilization of sucrose (Zacharaki *et al*., 2022). SUC2 gene in yeast is believed to function as invertase and mainly responsible for sucrose utilization (Singh *et al*., 2023). SNF1 is a positive regulator of SUC2 in yeast cells (Neigeborn and Carlson, 1984) and previous reports suggest that upon glucose depletion, SNF1 regulates the genes responsible for utilization of different carbon source, respiration and gluconeogenesis genes (Gancedo, 1998; Vincent and Carlson, 1999). Previous studies demonstrated that SNF1 is essential for glycogen accumulation (Cannon *et al*., 1994; Wiatrowski *et al*., 2004) and maintenance of glycogen store in the yeast cells (Wang *et al*., 2001). Further, we were curious to know the effect of TaSnRK1α in glycogen accumulation. As expected, both the qualitative and quantitative analysis show the more glycogen accumulation in Δ*snf1*+*TaSnRK1* complemented yeast cells as compared to Δ*snf1*. Moreover, there was not much comparable difference observed between CEN.PK and Δ*snf1*+*TaSnRK1* complemented cells. Histochemical assay clearly shows the difference in glycogen accumulation which was further supported by quantitative analysis. Furthermore, the stored glycogen showed the characteristic spectra by FTIR. The peaks at 1000 to 1200 cm-1 correspond to glycosidic bonds in glycogen and starch (Wood *et al*., 2004; Toepel *et al*., 2008). Overall, our results show the ability of TaSnRK1α to restore the growth defect of Δ*snf1* and promotes glycogen accumulation in yeast cells. Moreover, ATP-binding domain and UBA domain are essential for TaSnRK1α function as the truncated protein fails to complement Δ*snf1* function.

### TaSnRK1α showed dual localization and enhance starch accumulation in tobacco leaf

To explore the localization of TaSnRK1α, GFP fused TaSnRK1α was infiltrated in tobacco leaves. It was interesting to note that TaSnRK1α showed dual localization and found to be localized in nucleus as well as in cytoplasm in tobacco epidermal cells. 35S:*TaSnRK1*α:GFP was found to be localized in nucleus as well as in cytoplasm of tobacco epidermal cells. The localization of SnRK1 is still unclear and poorly understood. Some reports suggested nucleus localization of KIN10 in Arabidopsis (Shi *et al*., 2022). Similar report identified nucleus localization of SnRK1 in barley (Han *et al*., 2020). However, another study by Williams *et al*., (2014) and Zacharaki *et al*., (2022) and reported nucleus as well as cytoplasmic signal of KIN10 in Arabidopsis protoplast. Further, transient over-expression of TaSnRK1α in tobacco leaves led to more starch accumulation suggesting its function in starch metabolism in plants. So, we reported the dual signal of the TaSnRK1α and its role in high starch accumulation, although the exact mechanism of SnRK1 is not well characterized in tobacco.

### *TaSnRK1*α overexpression modulate starch accumulation in Arabidopsis

A diverse function of SnRK1α is well reported in stress, energy-sensing, developmental processes and hypocotyl elongation but its role in starch biosynthesis is not well characterized. However, in a study by Peixoto *et al*., (2021), O/E of SnRK1α1 resulted to significant increase starch content in Arabidopsis but the exact mechanism is not reported. When we overexpressed *TaSnRK1*α in wild type Col 0, high starch content was observed in *TaSnRK1*α O/E lines. The findings were consistent with the effect of overexpression of SnRK1 in potato resulting enhanced starch content by upregulating the activity of SuSy and AGPase (McKibbin *et al*., (2006). Also overexpression of *SnRK1* increases activity of AGPase and SSIII and increased starch content in transgenic tobacco Jiang *et al*., (2013) and sweet potato Wang *et al*., (2017). Surprisingly, the morphological analysis showed larger size starch granules *TaSnRK1*α O/E lines. Over-expression of *IbSnRK1* resulted to the change in starch morphology and enlarged starch granules in O/E lines in sweet potato Ren *et al*., (2019). The enlarged starch granules might contribute to an increase in total starch content and practically more feasible to extract for industrial application. Starch shows a characteristics distinct peak at around 1000 cm^-1^ contributed by the presence of glycosidic bond. The peak intensity is directly proportional to the more glycosidic bonds reflecting the more starch content. Considering this, our FTIR interpretation show more intense and sharper peak at 1000 cm^-1^ in case of *TaSnRK1*α O/E lines, indicating a high starch content.

Further Arabidopsis mutant *kin10*, a T-DNA insertion mutant, disrupted in KIN10 (catalytic sub-unit of SnRK1), was used to validate the effect of TaSnRK1α. Complementation with *TaSnRK1*α leads to restoration of starch measured at the end of day, indicating the probable role of SnRK1 in starch metabolism. Surprisingly, it was noteworthy that the starch granule morphology and FTIR spectra quietly resembles to the wild type Col-0, suggesting TaSnRK1α can complement the function of *kin10* mutant and have role in starch biosynthesis. Altogether, these findings, for the first time reveals the function of TaSnRK1α in enhancing the starch content and granular morphology.

Starch biosynthesis is a complex process, involving multiple genes, like AGPase, starch synthases, GBBS, SBE and other biosynthetic enzymes (Jeon *et al*., 2010). Notably, O/E of *TaSnRK1*α resulted in up-regulation of starch pathways genes, including starch synthases, GBSSI and SBE. Meanwhile, we did not find any significant change in AGPase expression level. However, SnRK1 overexpression is associated with increased AGPase activity. An increase of 30-92 % in AGPase activity is reported as a result of *IbSnRK*1 gene overexpression in tobacco (Jiang *et al*., 2013). Similar findings were reported in several other plants (Jossier *et al*., 2009; Wang *et al*., 2017; Ren *et al*., 2019) where the change in carbohydrate content could be the result of modification of AGPase activity.

### TaSnRK1α modulates AGPase activity in transgenic Arabidopsis

AGPase is a key and rate limiting step in the starch biosynthesis pathways (Goren *et al*., 2018). The positive correlation between AGPase LS and starch content is previously reported in wheat and rice (Kang *et al*., 2013; Sakulsingharoj *et al*., 2004). But the mechanism of regulation is not known. It was previously reported that AGPase activity is regulated at post translational level mainly by its redox activation (Hendriks *et al*., 2003; Tiessen *et al*., 2003; Jossier *et al*., 2009) However, recent report supports a scenario where AGPase undergoes phosphorylation by the protein kinases (Ferrero *et al*., 2020). But it is still unclear about the phosphorylating agent responsible for AGPase activation.

To advance our knowledge about the effect of TaSnRK1α kinase on AGPase activity, firstly we measured the AGPase enzymatic activity and as expected high AGPase activity was found in *TaSnRK1*α O/E lines in comparison to the Col-0. Moreover, the AGPase activity in *kin10*+*TasnRK1*α complemented lines was found to be similar as Col-0. Results of the present study are consistent with other plant studies where they show increased enzymatic activity of AGPase in tubers, leaves and storage roots of plants of potato by overexpressing the SnRK1(McKibbin *et al*., 2006; Jiang *et al*., 2013; Wang *et al*., 2017; Ren *et al*., 2019). Further to give more support to our finding, we performed the in-gel enzymatic activity assay (zymogram). The zymogram results clearly show the difference in AGPase enzymatic activity when supplied with Glucose-1-P and ATP as substrate. The higher enzymatic activity follows the order as *TaSnRK1*α O/E > *kin10*+*TaSnRK1*α > Col-0 > *kin10*. Our findings also support the scenario where activity of AGPase is modulated by its phosphorylation probably by TaSnRK1α in case of wheat and could contribute directly towards the high starch biosynthesis.

Previous characterization of posttranslational modification of AGPase enzyme suggest that large subunit could be the major target of phosphorylation (Ferrero *et al*., 2020). So, we investigated the interaction between the TaSnRK1α and AGPase LS using *in-vitro* and *in-planta* studies. To our surprise TaSnRK1α was found to interacts directly with AGPase LS as seen in yeast two hybrid assay which was further confirmed using BiFC assay. To date no direct interaction between AGPase and TaSnRK1α is reported. Our study provides the first glimpse of physical interaction between AGPase LS and TaSnRK1α. Phosphorylation dependent activation of AGPase by TaSnRK1α could prove a new approach to alter the activity of this rate-limiting enzyme. This post-translational modification of AGPase enzyme could enhance the specific AGPase activity which is followed by the active biosynthesis of starch, a major carbon reserve in the grain.

In conclusion, our results reveal for the first time, the role of TaSnRK1α in regulating starch biosynthesis in wheat. Its overexpression increases not only starch content in Arabidopsis but also change the starch granule morphology. We also demonstrated that TaSnRK1α interacts with AGPase LS thereby modulating its activity (Fig. 7). The overall results suggest the potential role of TaSnRK1α in improving starch content of plants in future.

**Fig. 7:**
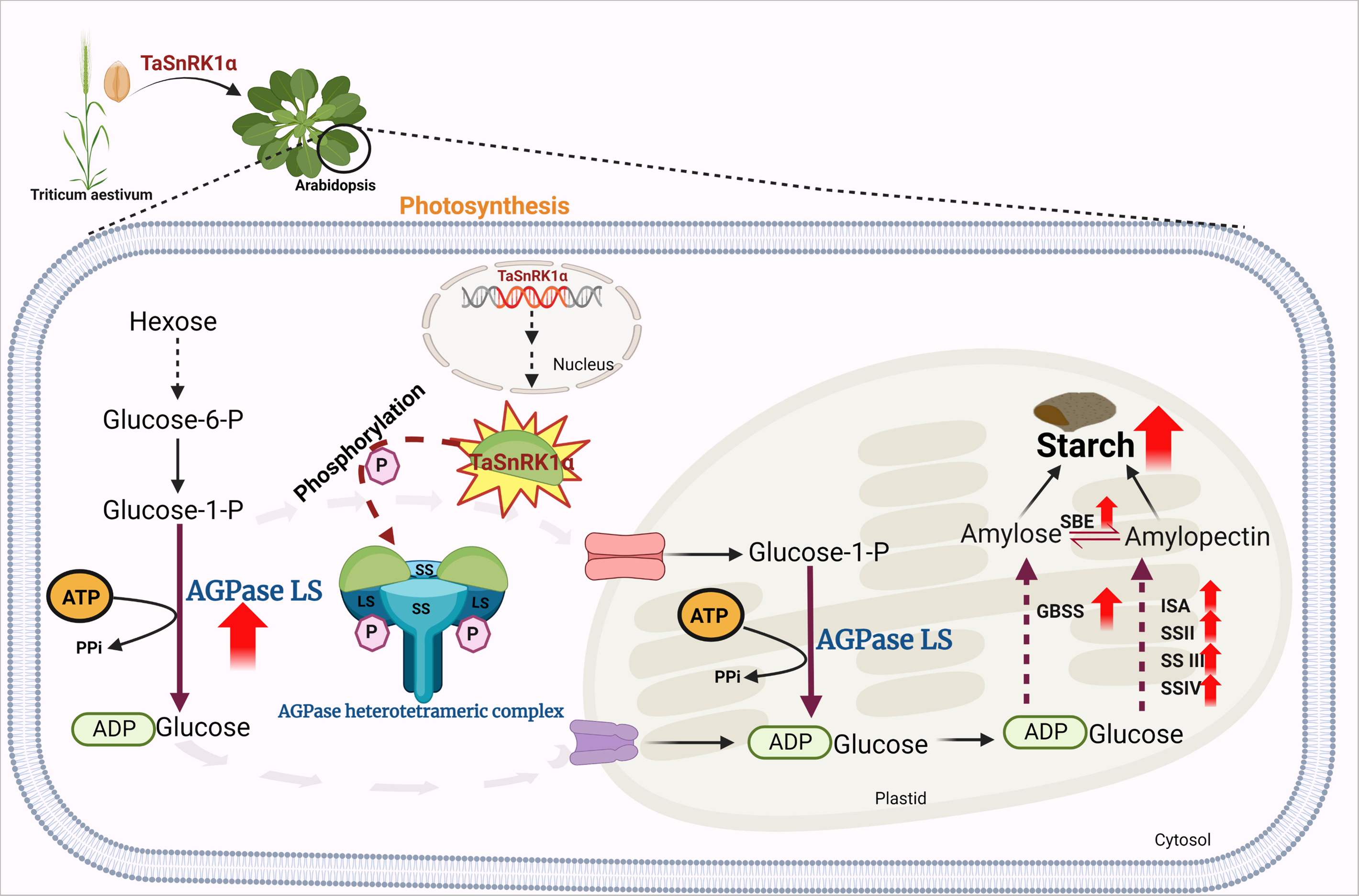
Diagrammatic representation of TaSnRK1α mediated regulation of starch biosynthesis pathway. Overexpression of TaSnRK1α increases AGPase activity. AGpase is a heterotetrameric complex with two large and two small sub-unit. Larger subunits are probably the target site for phosphorylation, thereby, activating the AGPase complex. The enhanced AGPase activity in turn contributes to more starch accumulation.

## Supplementary data

**Supplementary Fig. S1:** Light micrographs of yeast cells after iodine staining for glycogen content.

**Supplementary Fig. S2:** PCR confirmation of TaSnRK1α overexpression in Arabidopsis ecotype Col-0.

**Supplementary Fig. S3:** PCR confirmation of TaSnRK1α complemented in Arabidopsis KIN10 mutants.

**Supplementary Table 1:** List of primers used in the study.

## Acknowledgments

The authors are thankful to National Agri-Food Biotechnology Institute (NABI), an autonomous institute of Department of Biotechnology, Government of India (GOI) for providing research facility and financial support. We would like to thank Dr. Sunil Laxman (InStem, Bangalore) for providing wild type strain CEN.PK and yeast mutant Δ*snf1*. PK is also thankful to the University grant commission (UGC), India, for granting Junior and Senior Research Fellowship (JRF & SRF). PK also acknowledge Regional centre of biotechnology (RCB), Department of Biotechnology, Government of India (GOI) for undertaking PhD works. We also acknowledge DeLCON (DBT electronic library consortium), Gurugram, India, for providing access to e-resources.

## Author Contributions

PK and JKR conceived the research plan and designed the experiment. PK, AM and VS performed the experimental work and data analysis. AS and DS assisted in Zymogram assay. AP, DD and VF finalized the manuscript writing. PK, AM, VS, AS, DS, JKR edited and draft the final manuscript. All the authors commented, read, and approved the final manuscript.

## Conflict of interest

The authors declare no conflict of interest.

## Data availability

All the data supporting the findings of this study are available within the paper and its supplementary data published online. All raw data underlying the results presented in this study are available upon request.

## Abbreviations

ADP: Adenosine diphosphate
AGPase: ADP-glucose pyrophosphorylase
ATP: Adenosine triphosphate
AMPK: The AMP-activated protein kinase
SnRK1: SNF1- related protein kinase 1.

## Notes

### Competing Interest Statement

The authors have declared no competing interest.

